# TelOMpy enables single-molecule resolution of telomere length from optical genome mapping data

**DOI:** 10.1101/2025.03.11.642545

**Authors:** Ivan Pokrovac, Sunčica Stipoljev, Željka Pezer

## Abstract

The existing methods for assessing telomere length are limited in various ways, including resolution, scope, throughput, and reproducibility. These limitations stem from the applicability of laborious and complicated indirect approaches. We developed TelOMpy, a bioinformatics tool for determining the lengths of individual telomeres from optical genome mapping data. TelOMpy directly measures the absolute lengths of thousands of telomeres within a single sample, eliminating errors associated with indirect methodologies and ensuring high analytical power and reproducibility of results. We used optical genome mapping data generated from mice in a high-fat diet experiment to demonstrate the power of TelOMpy performance at the single-molecule level. Moreover, using our newly developed protocol for high-molecular-weight DNA extraction, we produced the first optical genome maps from sperm and compared telomere lengths between sperm and somatic tissue in overweight mice. TelOMpy reveals immense variation in the length of individual telomeres and the effect of a high-fat diet on telomere shortening and loss.

## Introduction

Telomeres are specialized structures on chromosome ends that ensure chromosome integrity. At the nucleotide level, they are formed by stretches of a tandemly repeated short DNA motif. The lengths of these stretches are variable and relevant for aging-related processes and diseases (Blackburn et al., 2015). Multiple methods are available for assessing telomere length. One of the most widely used techniques is quantitative PCR (qPCR; Cawthon, 2002), which requires a small quantity of DNA and a simple, inexpensive procedure for sample preparation. Specifically, the qPCR-based methods determine the average telomere length between DNA molecules present in the qPCR reaction relative to a reference sample. A major downside of this approach is the low reproducibility between studies (Aviv et al., 2011; Martin-Ruiz et al., 2015). Terminal restriction fragment (TRF) is another method for measuring average telomere length in a population of cells that is insensitive to very short telomeres (Montpetit et al., 2014). On the other hand, single-telomere length analysis (STELA) and quantitative fluorescence in situ hybridization (Q-FISH) are methods of choice for assessing the length of single telomeres with a resolution of 0.1 kilobase pairs (kbp). However, while the former is usually limited to telomeres of a few chromosomes, only cells in metaphase can be assayed with the latter (Aubert et al., 2012; Montpetit et al., 2014). While qPCR-based methods can be scaled up to screen hundreds of samples, essentially all these methods are low-throughput.

Optical genome mapping (OGM, Bionano Genomics) is a high-throughput genome analysis technology that employs imaging to construct whole-genome maps. High-molecular-weight DNA (HMW DNA) is fluorescently labeled via the DLE-1 enzyme on the CTTAAG motif, which is repeated in the human and mouse genomes on average every 10–15 kbp. Long DNA molecules are stretched through nanochannels under an electric field, and fluorescent signals are detected at labeled positions (Yuan et al., 2020). The imaged molecules are digitized, and the alignment of molecules is performed based on the pattern of DLE-1 motif positions (*labels*) to create consensus maps (CMAPs) of genomic regions. These maps can be further used to construct genomes *de novo* and to detect genomic structural variants on the basis of comparisons with the *in silico* labeled reference genome. As a single-molecule technique, OGM enables the direct study of individual DNA molecules without the need for fragmentation or amplification, which eliminates errors related to these procedures. However, given that the result of OGM are genome maps of labeled motifs, information on nucleotide sequences is not available from the obtained data.

We present TelOMpy, a bioinformatics tool that measures the absolute lengths of individual telomeres from optical genome mapping data. Within a single sample, TelOMpy directly calculates the lengths of thousands of telomeres from long, high-quality molecules aligned to chromosome ends of a reference genome. TelOMpy can be applied to any species for which chromosome-level genome assembly is available. We demonstrate its usage in mice in a high-fat experiment. Overweight and obesity are known to be associated with telomere shortening in somatic and reproductive cells (Valdes et al., 2005; Gielen et al., 2018; Clemente et al., 2019; Yang et al., 2016; Raee et al., 2023), and this association may have harmful consequences for offspring through maternal or paternal effects (Yang et al., 2015; Yang et al., 2016; Wei et al.; 2021). It is therefore of interest to study the effects of diet on telomere length in both somatic and reproductive cells. As part of this research, we developed a protocol for the extraction of high-molecular-weight (HMW) DNA from mouse sperm. This allowed us to perform the first OGM on sperm cells and apply TelOMpy to calculate the lengths of thousands of telomeres and to compare them with telomeres from the kidney as a representative of somatic cells. Our results illustrate the ability of TelOMpy to perform high-throughput measurements of telomere lengths at an unprecedented resolution, enabling the detection of subtle differences. Our study highlights the tremendous variation in telomere length between and within individuals and chromosomes and the effect of a high-fat diet on telomere shortening and loss.

## Results

### TelOMPy overview

Optical mapping is a method based on single molecules, which entails the capacity to determine the lengths of individual telomeres. These are calculated from the molecules that are mapped to the end label on a chromosome arm in the reference genome as the distance between it and the end of the molecule. This calculation of telomere length is based on the absence of a DLE-1 motif in telomeres, as they typically consist of a tandemly repeated TTAGGG motif. In addition to the telomeric sequence, such a measurement includes a part of the subtelomeric region with no DLE-1 motif. Hence, the length of the nontelomeric part can be calculated as the distance from the last aligned label on the reference genome to the start position of the annotated telomere (Figure 1).

**Figure 1.**
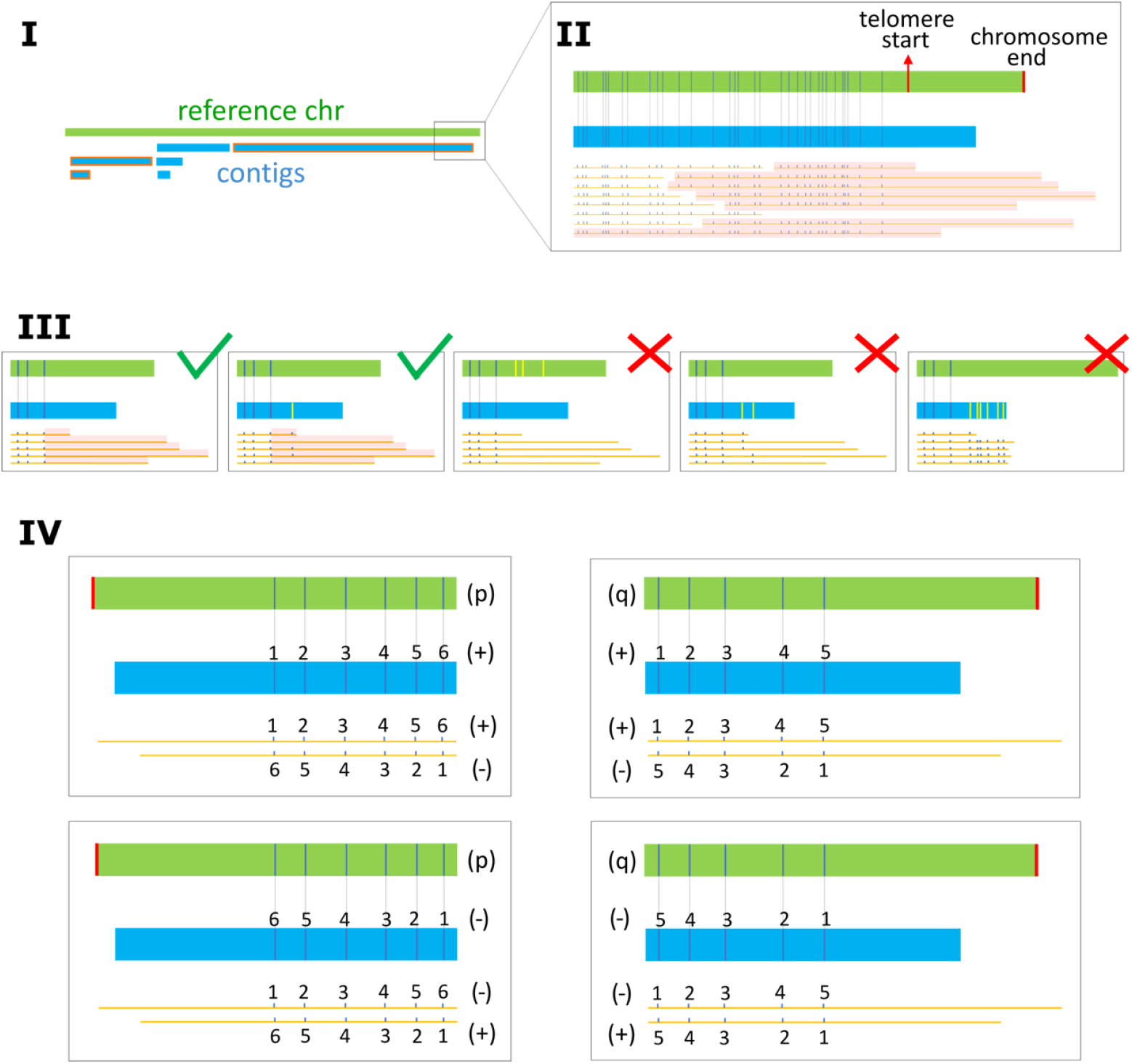
TelOMpy workflow. Throughout the figure, the reference chromosome is depicted in green, and the contigs are depicted in blue rectangles; the molecules are shown as orange horizontal lines with dark blue ticks as labels; the labels on the contigs and reference are shown as vertical dark blue (aligned) and yellow (unaligned) lines; and the alignments between the labels on the reference and contig are indicated by connecting light gray vertical lines. I) Contigs aligned to chromosome ends are identified (outlined in red) from the OGM *de novo* assembly. II) Telomeric molecules (highlighted in light red) are identified as molecules that are aligned with the telomere ends of these contigs. Only a long chromosome arm telomere is illustrated, but telomeres on both chromosome arms are considered for analysis by TelOMpy. III) Alignments on chromosome ends are checked, and only those of the highest quality (see Methods) are retained for analysis. Individual telomeres are highlighted in light red as part of the telomeric molecule after the outermost aligned label. IV) Telomere lengths are calculated by applying a proper equation to each telomeric molecule based on the chromosome arm and orientations of the molecule and contig. The most distal parts of the p-arm (left) and q-arm (right) are sketched with all possible combinations of orientations of molecules and contigs. The position of the chromosome end on the reference genome is shown by a red vertical line. The orientation of the molecule and contig is indicated by “+” and “-”signs and illustrated by ascending or descending label numbers in telomeric molecules.

TelOMpy takes as input the results of *de novo* genome assembly generated by the OGM and determines telomere length in four steps (Figure 1): I) it first identifies contig(s) aligned to the ends of a chromosome, and II) it identifies *telomeric molecules*, i.e., molecules that are aligned to the telomere ends of these contigs. Next, III) the alignments are checked for suitability by counting the number of the most distally positioned labels that are continuously unaligned and by measuring the distance between the annotated chromosome end and the closest label. Ideally, the most distal label on the reference chromosome should also be the last aligned label. One unaligned label position is tolerated on contigs or molecules to account for the possibility of unspecific labeling or mutations, which may create a DLE-1 motif in telomere repeats. In addition, the greater the distance between the chromosome end and the closest label on the reference genome is, the more unlikely it is to be bridged by the molecules in the OGM. Therefore, in such cases, the most distally aligned molecules are not those of telomeres but rather represent the last genomic region where alignment was still possible, adjacent to a genomic “desert” without labels. Chromosome arms that are suitable for analysis are retained, and in the last step, IV) their telomere lengths are calculated by applying one of the two equations on every individual telomeric molecule, depending on the chromosome arm and the orientations of the alignments (see Methods).

### Newly established protocol yields high-quality DNA from sperm cells suitable for optical genome mapping

To test the performance of TelOMpy and further examine the link between overweight/obesity and telomere length in somatic and sperm cells, we performed a high-fat diet experiment on male mice. At a prepubertal age, the mice were fed a diet rich in fat, cholesterol, and sugar (Western diet group, WD) for 80 days. In parallel, mice of the same age were maintained on a standard diet throughout the entire experiment (control group, C). Although there was no significant difference in weight between the two groups at the beginning of the experiment (Mann-Whitney test, p value 0.58), mice in the WD group were expectedly heavier at the end of the experiment (Mann-Whitney test, p value 0.0086, Cohen’s D of 1.9, CLES 0.97), indicating the effect of a high-fat diet (Figure 2A).

**Figure 2.**
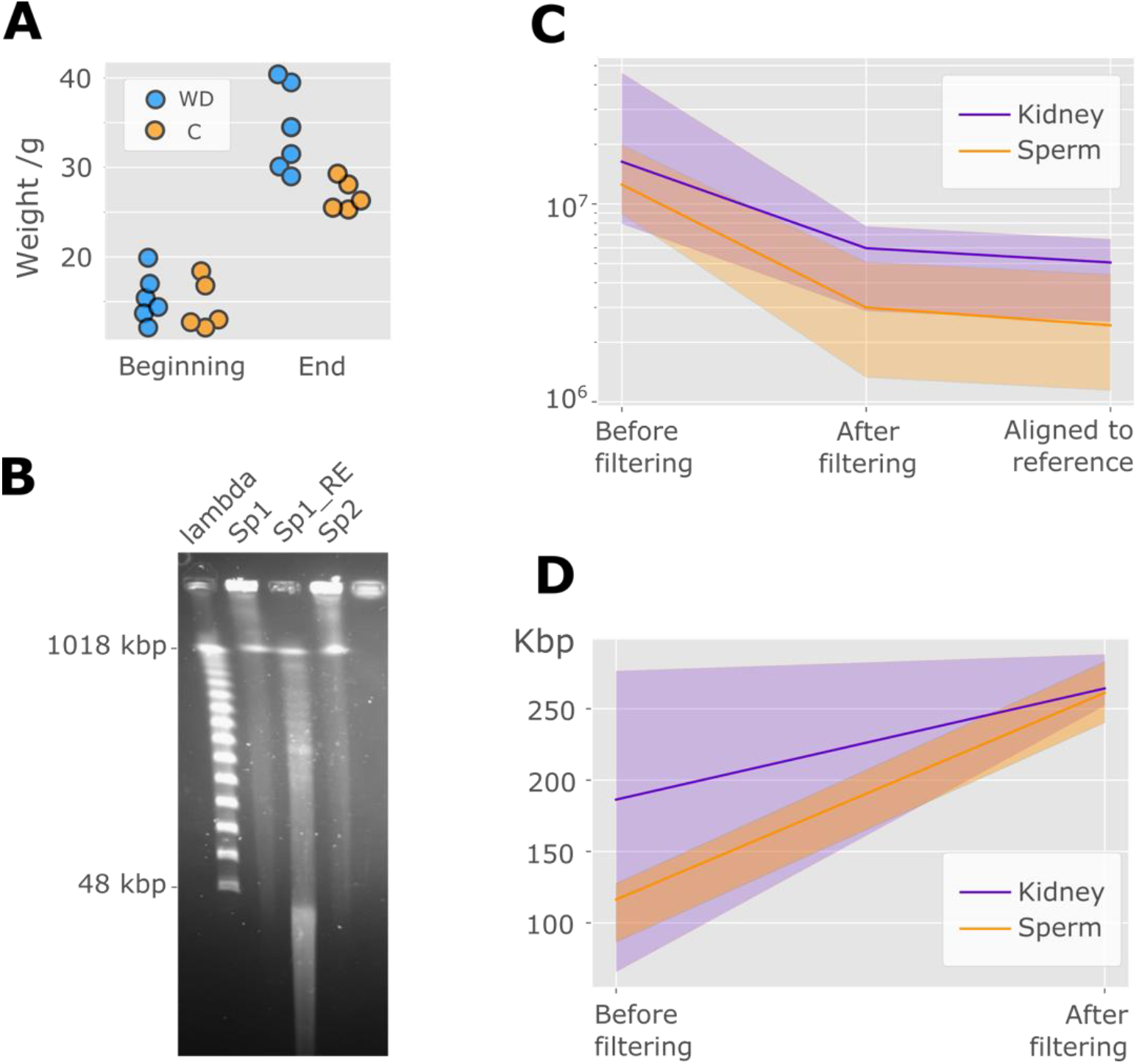
DNA and data quality indicators. A) Weights of the mice at the beginning and end of the experiment. B) An example of PFGE results. Lambda is a PFG ladder with a fragment range from 48.5 kbp to 1018 kbp. Lanes labeled *Sp1* and *Sp2* contain HMW DNA isolated from approximately 2.5 million sperm cells each. Lane *Sp1_RE* contains *Sp1* digested with the EcoR1 enzyme. C) Number of molecules produced by OGM at various stages of the *de novo* genome assembly process. D) Average length of molecules before and after filtering. In both C) and D), the lines represent the mean values, and shading shows the range of values between samples.

Kidney tissue and seminal fluid were dissected from each mouse. High-molecular-weight (HMW) DNA was extracted from kidney cells via the standard Bionano protocol (see Methods). For the extraction of HMW DNA from sperm cells, we modified the procedure by adding dithiothreitol (DTT), which acts as a reducing agent by breaking the disulfide bonds in protamines, proteins that are specifically present in sperm chromatin (Balhorn, 2007). A portion of the isolated HMW DNA was digested with a restriction enzyme to establish the efficiency of protein removal. The success of digestion and the integrity of the isolated DNA were checked via pulsed-field gel electrophoresis (PFGE). The size of the DNA before digestion was mainly greater than 1 Mbp (Figure 2B). After digestion, the length of the DNA did not exceed 1 Mbp, and a significant amount was less than 48 kbp. We concluded that the HMW DNA obtained from sperm cells with the established protocol is in the size range that meets the requirements for OGM (minimum size of 150 kb) and is free from proteins; thus, it is accessible for labeling with the DLE-1 enzyme.

Optical genome mapping was performed on HMW DNA isolated from the kidney and sperm cells of five mice in the experimental group and four mice in the control group. The DNA concentration after DLS labeling ranged within Bionano’s recommendations (Table S1, Supplementary Material).

### Quality of optical genome mapping data

To establish how the newly developed protocol for HMW DNA extraction affects the quality of molecules and *de novo* genome assembly, we compared the data between sperm and kidney samples. Multiple metrics indicate the high quality of the resulting OGM data after *de novo* assembly (Table S2, Supplementary Material). All samples meet the recommended quality criteria for the average length of molecules, effective coverage of the reference, and fraction of molecules aligned. Moreover, in all the samples, the N50 of the diploid genome map was much greater than the recommended 50 Mbp minimum, indicating high continuity of the assembly.

DNA molecules that enter the OGM are filtered during the *de novo* genome assembly pipeline: those shorter than 150 kbp and containing fewer than 9 labels are considered low-quality and are thus discarded. We compared the number of molecules before and after the removal of low-quality molecules. Although the initial number of molecules is similar in both tissues, more molecules are filtered out in the sperm samples, resulting in approximately three times more molecules aligned to the reference genome in the kidney than in the sperm samples (Figure 2C). The more extensive removal of initial molecules in the sperm samples resulted in differences in the effective coverage of the reference genome, which was significantly lower for the sperm samples than for the kidney samples (paired t test, p value 0.0007, Cohen’s D 2.53); however, this coverage was still above the recommended coverage minimum (Table S2, Supplementary Material). Although the average length of molecules isolated from sperm was shorter than that from kidneys, this difference disappeared after the removal of low-quality molecules (Figure 2D).

In conclusion, compared with the standard Bionano protocol for HMW DNA isolation from somatic tissue, the newly developed procedure for isolation from sperm initially yields lower-quality molecules on average; however, after filtering, the remaining number of molecules is sufficient to achieve high-quality *de novo* genome assembly.

### Data suitability

TelOMpy was applied to OGM data generated from the kidneys and sperm of mice in the Western diet experiment. As described above and in the Methods section, information on telomere length is only available from high-quality alignments of molecules and contigs with the reference genome at chromosome ends (Figure 1). Owing to this requirement, it was not possible to determine the telomere lengths on the short (p) arms of almost every chromosome in all the samples (Figure 3A). Mouse chromosomes are acrocentric, containing regions of unknown sequence and length between the telomere on the p-arm and the centromere. These regions are annotated in the reference genome as gaps of an arbitrarily chosen length of 10 kbp. In our samples, they were devoid of labels corresponding to the DLE-1 enzyme, and the molecules obtained by OGM were not long enough to bridge the p-arm telomere and centromere regions and provide information on telomere length. We encountered a similar problem on the long (q) arms of the sex chromosomes and chromosome 4. Additionally, chromosomes 9 and 14 of some samples had one or more unaligned labels on the reference after the last aligned label (Figure 3A). Finally, for these reasons, all short chromosome arms as well as long arms of sex chromosomes and chromosomes 4, 9, and 14 were excluded from subsequent analyses of telomere length in all samples. We extracted telomeric molecules aligned to the remaining chromosome arms. To estimate the degree of their alignment in subtelomeric regions, we calculated the proportion of molecules with different numbers of unaligned labels at distal positions. Approximately 90% of telomeric molecules support contigs whose outermost label is aligned with the outermost label on the reference genome, and approximately 10% are aligned to contigs with a single unaligned label at the most distal position. In more than 80% of all telomeric molecules, the outermost label was aligned with the outermost label on the reference, and approximately 98% had fewer than four continuously unaligned labels at distal positions (Figure 3B). These data suggest that the degree of alignment of molecules at the end of the q-arm of most chromosomes is suitable for telomere length analyses.

**Figure 3.**
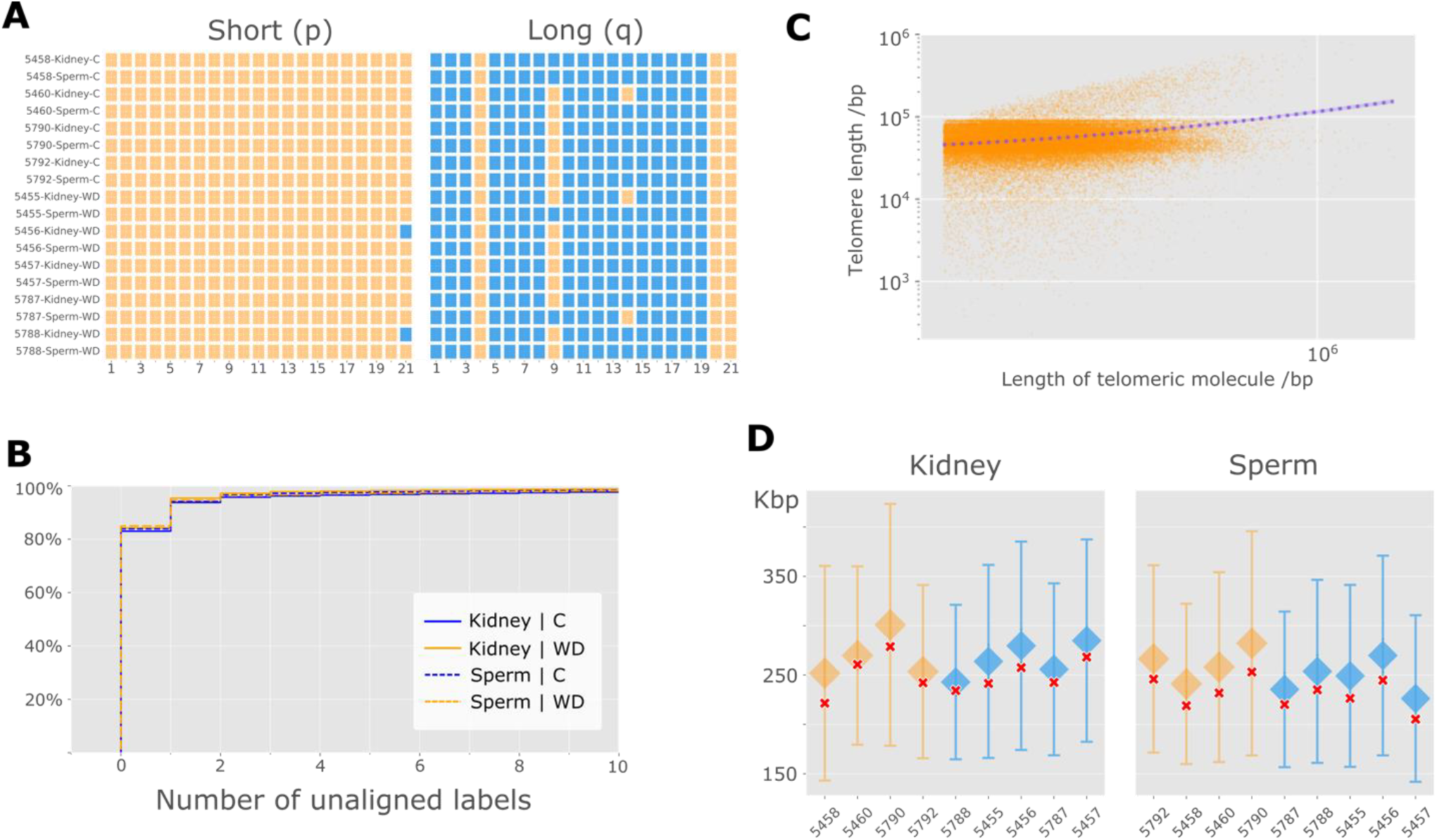
Data suitability for the analysis of telomere length. A) Overview of chromosome arms suitable for telomere length determination by sample. Chromosome arms at which alignments do not meet the quality criteria are marked in orange, and chromosome arms at which alignments are suitable for analysis are marked in blue. Information for autosomes 1--19 and sex chromosomes (designated 20 and 21) is shown in columns for each sample per row on the p and q chromosome arms. B) Cumulative empirical distribution for the number of continuously unaligned labels at distal positions on the molecule. C) Correlation between the length of molecules carrying telomeres and the calculated telomere length. D) Distribution of the length of telomeric molecules per mouse in the kidney (left) and sperm (right) for the control (orange) and experimental groups (blue). Mouse identification numbers are shown on the x-axis. The diamond shapes indicate the average length of telomeric molecules, and the caps indicate one standard deviation of the average length. The red sign x indicates the average length of all molecules aligned to the reference genome.

The calculated lengths of telomeres in our dataset may not reflect true lengths but rather shorter artifacts that stem from degradation processes that may have occurred at chromosome ends during cell lysis. To control for this possibility, we compared the lengths of telomeric molecules with the lengths of all the molecules that entered the alignment process. We found no significant correlation between telomere length and the length of molecules from which telomere length was calculated (Pearson’s R=0.24; Figure 3C). Similarly, we found no significant difference between the length of molecules carrying telomeres and the average length of all molecules aligned to the reference genome (Cohen’s D < 0.3; Figure 3D). These results indicate that the molecule length in our dataset does not have a significant effect on the determination of telomere length.

Considering that the telomere length from the optical mapping data is calculated from the last DLE-1 motif to the end of the molecule, each measured telomere length also contains a part of the subtelomeric region between the last motif DLE-1 and the beginning of the annotated telomere on the reference genome. This part varies between chromosomes on the mouse reference genome and amounts to tens to thousands of base pairs (Supplementary Figure S1) and can be considered negligible compared with the average mouse telomere length of 50 kbp (Hemann, 2000). These values can be subtracted from the calculated telomere lengths to obtain corrected absolute lengths. However, their inclusion does not affect the comparison of telomere lengths between samples since this factor is equally present in calculations for all samples.

### Differences between the experimental and control groups

A total of 70,215 telomeres were compared in our dataset, with an average of 3,900 telomeres per sample (range 1,179–7,869). On average, we found a threefold lower number of telomeres in the sperm than in the kidneys. This can be only partially explained by the overall lower genome coverage in sperm samples, given that normalization for the number of aligned molecules only somewhat reduces the difference (Figure 4A). In addition, within both tissues, a smaller number of telomeres were detected in the mice in the experimental group. This difference is even more pronounced when we normalize the number of telomeres by dividing it by the total number of molecules. These results suggest that the lower telomere count in the WD group is a result of the experimental effect rather than a technical artifact (Figure 4A).

**Figure 4.**
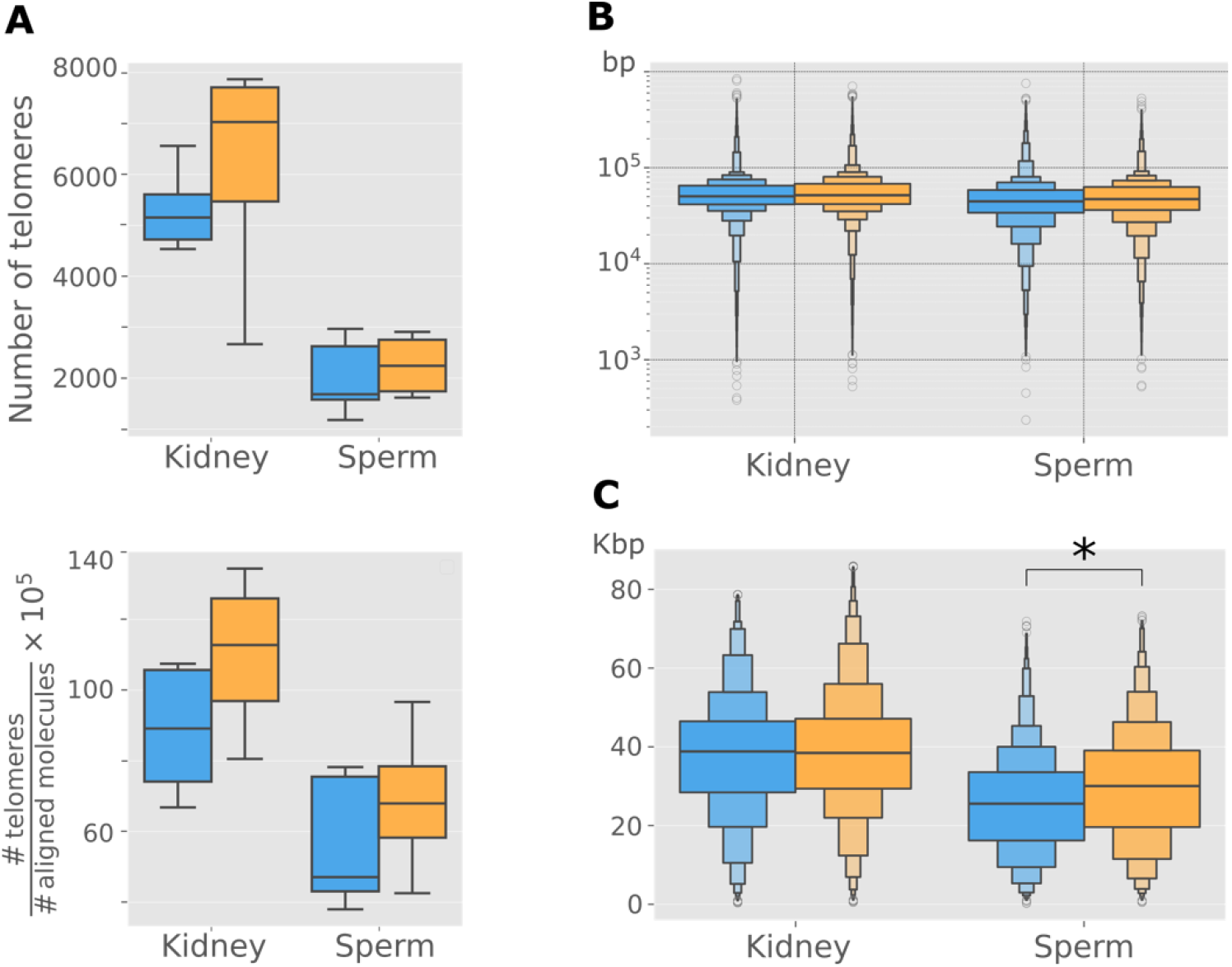
Comparison of telomere length between the experimental and control groups. Throughout the figure, the WD group is shown in blue, and the C group is shown in orange. A) Telomere count (top) and the normalized number of telomeres (bottom). This normalized number corresponds to the number of telomeric molecules per 100,000 molecules aligned in the genome. B) Distribution of telomere lengths shown by a “boxen” diagram by tissue. C) Distribution of telomere lengths in Q1. The horizontal line in the widest part of each of the distributions shown in B) and C) represents the median value.

The measured length of individual telomeres varies between 236 bp and 840 kbp, with an average of 55 kbp in the dataset. Telomeres are on average shorter in mice from the experimental group (Figure 4B), by 2.6 kbp (median 1.3 kbp) in kidney tissue (t test, p value 4 × 10^−25^) and by 1.2 kbp (median 2.5 kbp) in sperm (t test, p value 4 × 10^−19^). However, given the large variation, this effect is small for both kidney tissue (Cohen’s D = 0.08) and sperm (Cohen’s D = 0.04). Despite the wide range of telomere lengths within the same chromosome, we observed significant differences in average telomere length between chromosomes, with 43 kbp being the shortest length on chromosome 10 and 74 kbp being the longest length on chromosome 11 (Figure S2).

To determine whether a high-fat diet induces specific telomere shortening in sperm cells, we compared the distribution of telomere lengths from sperm with the distribution of telomere lengths from the kidneys of the same individual. The distribution is shifted toward lower values in sperm than in kidneys in all individuals, which indicates a general shortening of telomeres in sperm and/or their lengthening in the kidney in both groups. There was no significant difference in this effect between the experimental and control groups: in the control group (N=4), the average length difference was 7.8 ± 4.4 kbp (median difference of 6.8 ± 4.2 kbp), and in the experimental group (N=5), it was 6.1 ± 3.1 kbp (median difference of 6.0 ± 2.8 kbp) (Figure S3).

Critically short telomeres can affect the replicative potential of the cell and cause genomic instability, apoptosis, premature aging, and a shorter lifespan (Hemann et al., 2001). Hence, while the average telomere length may not differ significantly, the shortest telomeres may cause dysfunction and are thus highly relevant for the study of telomere length. To analyze in more detail the differences in the length of critically short telomeres, we extracted the first 25% of the shortest telomeres (first quartile of length, Q1) within each chromosome of each individual to account for the significant variation in telomere length between individuals and between chromosomes. Within the first quartile of length, telomeres are shorter in the sperm of the experimental group than in those of the control group (t test, p value 9 × 10^−26^; Cohen’s D = 0.31) by 4.3 kbp on average (median 4.4 kbp). This difference in kidney tissue amounts to 1 kb and is not statistically significant (Table 1, Figure 4C). The difference in average telomere length in the first quartile of length between kidney tissue and sperm in the control group (N=4) was 10.1 ± 7.5 kbp (median 10.0 ± 7.2 kbp), and that in the experimental group (N=5) was 12.1 ± 6.6 kbp (median 12.5 ± 7.4 kbp) (Figure S4).

**Table 1.** Descriptive statistics of telomere length (in Kbp).

| Group | Tissue | Average | Average STDEV** | Median | Average IQR** |
| --- | --- | --- | --- | --- | --- |
| C | Kidney | 38.7 | 16.0 | 38.4 | 17.8 |
| WD | Kidney | 37.7 | 15.5 | 38.8 | 18.1 |
| C | Sperm | 30.0 * | 14.9 | 30.0 | 19.4 |
| WD | Sperm | 25.7 * | 13.1 | 25.6 | 17.4 |

\* Statistically significant difference between WD and C (Mann–Whitney test, P value < 0.05; Cohen D > 0.3)
\*\* IQR - interquartile range; STDEV - standard deviation

### Validation of TelOMpy-calculated telomere lengths

The wide range of telomere sizes and the average length determined by TelOMpy in our dataset are consistent with previous studies that used other methods to show comparable average and variation in telomere lengths in the C57BL/6 strain (Zijlmans et al. 1997; Hemann, 2000; Zhang et al., 2025), indicating the general reliability of TelOMpy.

To further validate our results obtained via TelOMpy, we performed a qPCR assessment of telomere length (Cawthon, 2002). For this purpose, a commercial kit was used to extract genomic DNA from samples of kidney and sperm from the same samples of mice in the described high-fat diet experiment. For each biological sample, the isolated genomic DNA was used as a template for singleplex qPCR, with primer pairs specific to either the telomere (T) or the albumin gene used as single-copy reference gene (S). Relative telomere length was expressed as the ratio of telomere copy number to the reference gene number (T/S). Our qPCR experiments confirmed shorter telomeres in the sperm than in the kidney (two-tailed paired t test; p value = 4.2 × 10^−3^) and no difference in telomere length between the control and experimental groups (two-tailed unpaired t test; p value = 0.785) (Figure 5). These findings were confirmed via qPCR on an extended dataset of kidney and sperm samples from 12 additional mice (see Methods and Figure S5 in the supplementary data).

**Figure 5.**
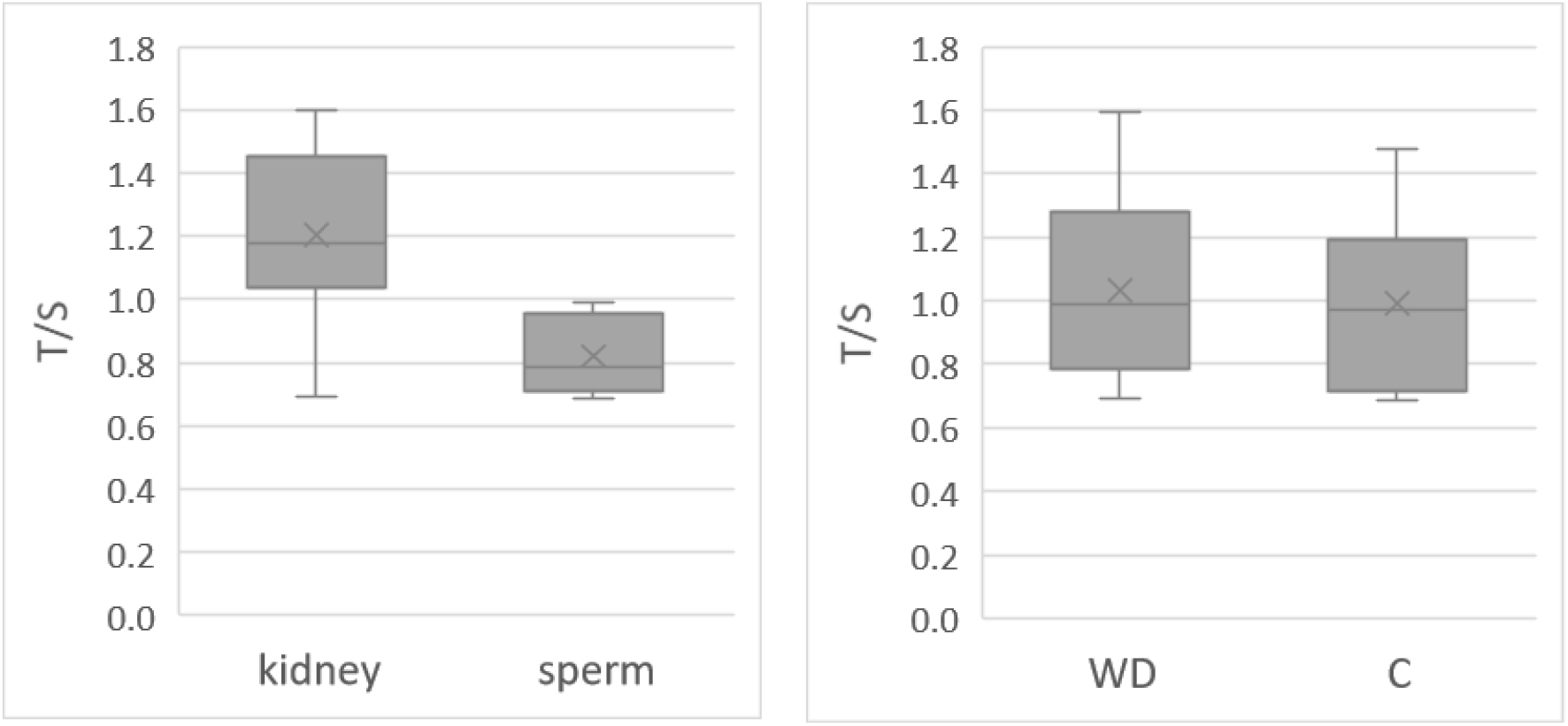
Validation of the TelOMpy results via qPCR. Telomere lengths relative to a control sample are presented as the T/S ratio in comparisons between tissues (left) and groups (right). N=8 for each of the plotted groups.

## Discussion

### Telomere length at the single-molecule level

Studying telomeres at the single-molecule level may provide important insights into which commonly used bulk approaches are blind. Recently, a tool called Telogator has been developed for the use of data produced via long-read sequencing technologies (Stephens et al. 2022). While this tool reports the lengths of individual chromosomes, it is limited by read length and thus is applicable only to species whose average telomere length does not exceed 10 kbp. Two methods for determining individual telomere lengths from optical mapping data have been described thus far. One of them requires the CRISPR/Cas9 system to mark the telomeric motif, which creates additional complications and costs in implementation (McCaffrey et al., 2017). The other has not been implemented in publicly available software (Young et al., 2017). Unlike the abovementioned methods, TelOMpy is a publicly available tool for determining a wide range of telomere lengths from optical genome maps, which does not require the use of additional technology. TelOMpy can be used at no extra cost in any research involving standard *de novo* genome assembly procedures and structural variant detection from OGM data. For example, OGM is often used to investigate genomic rearrangements in tumor tissues, in which changes in telomere length are part of cancerous processes (Fan et al., 2021).

We demonstrate the use of TelOMpy in a high-fat diet experiment in mice on OGM data produced from the kidney and sperm. We find that telomeres vary in length over three orders of magnitude—from only a few hundred to several hundred thousand base pairs. Such variability exists between and within individuals and tissues and between and within chromosomes and is consistent with telomere length polymorphisms previously demonstrated in human somatic cells (Lansdorp et al., 1996; Schmidt et al., 2024). Our results show that, on average, sperm telomeres are significantly shorter in the experimental group than in the control group, which is the detection power at the level of the qPCR measurement of telomere length. This result suggests that a high-fat diet is associated with telomere shortening in sperm. However, TelOMpy enabled more detailed comparisons at the level of individual chromosomes between kidney and sperm within a single mouse, which do not support this conclusion because no consistent shortening of telomeres exists exclusively in the sperm of experimental mice. These results demonstrate the unparalleled power and resolution of TelOMpy in analyses of telomere length.

Numerous studies have reported longer telomeres in sperm than in somatic cells due to the activity of telomerase in spermatogenic cells, which lengthens telomeres (Fice and Robaire, 2019). However, in all the analyzed mice in our study, telomeres were consistently shorter in the sperm than in the kidney. Our control analyses suggest that these results are not technical artifacts but rather represent a true biological state. We further confirmed this finding via qPCR on an extended sample set via a different method for genomic DNA extraction. However, telomere shortening in sperm has rarely been reported as a result of aging (de Frutos et al., 2016) or infertility (Tahamtan et al., 2019). In addition, mutations in various genes can lead to a decrease in telomerase activity and consequent telomere shortening in sperm (Margalef et al., 2018; Mirabello et al., 2010; Soerensen et al., 2012; Bojesen et al., 2013; Grill and Nandakumar, 2021). The relatively shorter telomere length in the sperm samples than in the kidney samples may be a result of mutations in one or more genes among the many involved in telomere maintenance. Since OGM data do not provide information on nucleotide sequences, it was not possible to perform analyses at the sequence level. Nonetheless, mutations that diminish or prevent telomerase activity are possible and may not necessarily be detrimental, as telomerase-deficient mice can be viable, fertile, and without visible morphological abnormalities (Blasco et al., 1997). Alternatively, the relatively longer telomeres in kidney cells than in sperm could result from telomerase activity in kidney cells. Mice are known to express telomerase in some somatic tissues (Prowse and Greider, 1995), and telomerase activity in various somatic tissues, including the kidney, has also been reported in chickens (Venkatesan and Price, 1998).

Surprisingly, we found that in both analyzed tissues, the number of telomeres was significantly lower in the experimental group than in the control group, indicating the loss of entire telomeres due to a high-fat diet. The complete loss of telomeres on chromosomes is a phenomenon previously reported in tumor cells (Fouladi et al., 2000; Lo et al., 2002; Lin et al., 2010; Raseley et al., 2023) and mutants (Vannier et al., 2012). Severely truncated telomeres are also common in normal cells of healthy humans, including sperm cells (Baird et al., 2003; Baird et al., 2006; Britt-Compton et al., 2006; Capper et al., 2007). Although a firm confirmation of our findings would require validation via alternative methods, our results suggest that diet can also induce complete telomere loss in both somatic and reproductive cells.

### Optical genome mapping of sperm cells

As part of this study, a protocol was developed for the isolation of HMW DNA from sperm for the purpose of optical genome mapping. The existing commercial protocol for the isolation of HMW DNA (Bionano Genomics) is optimized for somatic cells and is inefficient in breaking down the protein component of sperm chromatin, which is a crucial requirement for successful DNA labeling in OGM. We developed a procedure in which chromatin proteins are efficiently removed from the sperm genome while long HMW DNA molecules are isolated. This enabled us, for the very first time, to apply OGM technology to sperm genomes. We show that the developed protocol successfully yields high-molecular-weight (HMW) DNA of sufficient amount and quality and results in high-quality OGM data that are comparable to those of somatic tissue.

The OGM is widely used for detecting large structural variants (SVs). The high-molecular-weight DNA extraction protocol we present here facilitates the study of structural variation in sperm genomes, which represent the genetic material directly passed on to the next generation. In the OGM of sperm genomes, each DNA molecule originates from a single sperm cell, ensuring that each cell reflects a single allele of a structural variant at a specific locus. This approach enables two significant advancements. First, it allows the direct detection of *de novo* SVs in the germline, examining a population of sperm cells from an individual. Second, it enables the tracking of shifts in the frequency of existing variants within a single generation. These advancements create the potential to study the environmental impacts on structural variations, which can have direct consequences for the offspring phenotype.

## Methods

### High-fat diet experiment

Before the experiment, the mice were kept in groups of three mice per cage under standard laboratory conditions in the mouse facility at the Ruđer Bošković Institute in Zagreb, Croatia. Males of the C57BL/6 strain aged 4–5 weeks were randomly selected for the experiment. Six mice in the control group were fed a standard laboratory diet (4RF 21 Mucedola srl, Italy). Six mice in the experimental group were fed a Western diet (WD, ssniff EF R/M acc TD88137 mod, catalog # E15721-34, ssniff Spezialdiaten GmbH, Germany), characterized by high fat (21.2%), sugar (33.2%), and cholesterol (2.071 mg/kg) contents. Food and water were provided *ad libitum* in both groups. The experiment lasted 80 days.

In an extended experiment, six additional males of the C57BL/6 strain, aged 8–9 weeks, were randomly selected to repeat the high-fat diet experiment as described above. Six males of the same age were kept as a control group under standard conditions in parallel. Mice from the extended experiment were used solely for validation of telomere length via qPCR.

### Tissue preparation

One mouse from the control group died before the end of the experiment. At the end of the experiment, the remaining mice were euthanized by cervical dislocation. The kidneys and *cauda epididymides* were isolated from each mouse. Each kidney was cut into 2–3 pieces, corresponding to ~30 mg per piece, frozen by immersion in liquid nitrogen, and stored at −80 °C. *The cauda epididymis* was separated from the testis and transferred to an Eppendorf tube with 150 µL of DPBS buffer (Dulbecco′s phosphate-buffered saline, Sigma-Aldrich, catalog # D8537) for brief storage before further processing. *The cauda epididymis* was placed on a parafilm sheet and cleaned of adipose tissue. *The epidermis* was punctured approximately ten times with a needle (16--21G), and after the viscous contents were drained from it, it was transferred back into an Eppendorf tube with 1,000 µL of DPBS, placed horizontally in the hybridization oven and incubated for 1 hour at 37 °C with 6 rotations per minute (rpm). After incubation, the suspension was filtered through a 40 µm filter. Another 500 µL of DPBS was added to the tube and filtered again. The filtrates were collected and combined. A total of 10 µL of the combined filtrate was transferred to a Neubar chamber for sperm cell counting. The remainder of the filtrate was centrifuged for 10 minutes at 1,000 × g at 4 °C, and the supernatant was discarded by careful pipetting. The remaining pellet in the Eppendorf tube was immersed directly in liquid nitrogen and stored at −80 °C until DNA isolation.

### Isolation of high-molecular-weight (HMW) DNA and shipment

Approximately 30 mg of kidney tissue per mouse was placed in a cryotube and shipped on dry ice to the Institute of Applied Biotechnologies a.s. (IAB, Olomouc, Czech Republic). Isolation of HMW DNA from kidney tissue was performed by IAB via the *Bionano Prep SP Tissue and Tumor DNA Isolation Protocol* (Document Number: 30339).

Isolation of HMW DNA from sperm was performed according to the protocol in the Supplementary Data (Text S1). Briefly, 3–4 million sperm cells were enclosed per agarose plug before cell lysis, and dithiothreitol (DTT) was used as a reducing agent during lysis to break down the protein component of the chromatin. A portion of the DNA was digested with a restriction enzyme and compared with undigested DNA via PFGE to assess the accessibility and integrity of the isolated DNA (Supplementary Text S1). Each agarose block with HMW DNA isolated from sperm was inserted into a separate 5 mL tube. The tube was filled with wash buffer (10 mM Tris-HCl, 50 mM EDTA), closed, sealed with parafilm, and shipped at room temperature to IAB for optical genome mapping.

### Optical genome mapping

Optical genome mapping was performed on the Saphyr platform (Bionano Genomics). Genome optical maps were generated for a total of 18 samples from nine mice: five from the experimental group and four from the control group. For each mouse, OGM was performed on HMW DNA isolated from sperm and kidney tissue. Enzymatic labeling of HMW DNA from kidneys and sperm was performed according to the BNG protocol described in the document *Bionano Prep Direct Label and Stain (DLS) Protocol* (Document Number: 30206). The DNA concentration after DLS labeling was taken as a measure of initial DNA quality, and the coefficient of variation of the DNA concentration was calculated from three measurements. Initial data processing, including *de novo* assembly of the genome and detection of SVs, was performed by the IAB using the set of bioinformatics tools BioNano Solve, according to the BNG *Guidelines for Running Bionano Solve on the Command Line* (Document Number: 30205).

### TelOMpy workflow

To extract molecules from OGM data that contain telomeres, TelOMpy first identifies contigs that are aligned to the ends of chromosomes. This information is in the “exp_refineFinal1_merged.xmap” file under the output of the *de novo* assembly pipeline results (directory/output/annotation). By using the contig identification number (ContigID), information about the alignment of molecules to the specified contig is extracted from the XMAP file labeled “EXP_REFINEFINAL_contig{X}.xmap”, where X is the unique ContigID for the given dataset. Similarly, the same ContigID is used on corresponding CMAP files labeled “EXP_REFINEFINAL_contig{X}_q.cmap” to extract information on the molecule length and label positions of the aligned molecules. Depending on the chromosome arm, orientation of alignment between the contig and the reference chromosome, and orientation of alignment between the molecule and contig, one of eight different ways of calculating telomere length can be applied (Figure 1) and can be reduced to two equations:

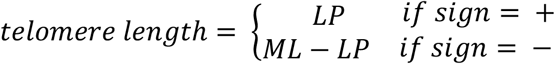

where *LP* represents the label position on the molecule that is aligned to the first (p-arm) or last (q-arm) label on the chromosome of the reference genome and where *ML* represents the total length of the molecule. If we assign a value of −1 to the long chromosome arm (q-arm) and a value of 1 to the short chromosome arm (p-arm), the *sign* in the above equation is defined as:

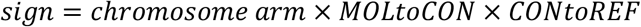

where *MOLtoCON* refers to the alignment orientation of a molecule to a contig and *CONtoREF* refers to the alignment orientation of a contig to a reference chromosome.

Cases where the most distal label on the reference genome is not aligned (paired) are controlled through the parameter *number of unpaired labels on the reference*. In the default settings of the TelOMpy tool, the value of this parameter is 0. However, this parameter can be set to values greater than zero if the distance to the most proximally aligned label on the reference is short enough for a molecule to fully encompass a telomere. The user is directed to check such cases visually in the Bionano Access browser and to account for possible species specificity in telomere length, label density, and average molecule length.

Similarly, unaligned labels on contigs or molecules, which are distant from the last aligned label on the reference genome, are controlled by introducing the parameters *number of unpaired labels on the molecule* and the *number of unpaired labels on the contig*. Some molecules or contigs may have one unaligned label in telomeric regions due to mutation or nonspecific labeling. Importantly, these parameters refer to the number of continuously unaligned (*unpaired*) labels to the left (on the p-arm) or the right (on the q-arm) from the outermost aligned label on the reference genome and should not be confused with the total number of unaligned labels on a molecule or contig, which may or may not be higher.

If the outermost label on the reference chromosome is aligned with a label on a contig but its distance from the annotated chromosome end (in the reference genome) is significantly greater than the average length of molecules, it is unlikely that the most distally aligned molecules will contain telomeres. For example, the first label position on chromosome 1 of the mm10 assembly is located 3 Mbp after the chromosome start position. To control for this, we introduced the *distance from the end* parameter as the maximum distance (in nucleotide base pairs) between the first/last aligned label and the end of a reference chromosome. This value, in addition to depending on the average length of the molecules, also depends on the length of the annotated telomeres for a particular organism. Telomeres are annotated on the reference genome as gaps of an arbitrarily set size, which is 100 kbp in mice and 10 kbp in humans (UCSC Table Browser), and the user should take this into account when defining the value of the *distance from the end* parameter. In the case of the mouse mm10 genome assembly, a value of 200,000 (200 kbp) was used, which is twice the length of annotated telomeres but less than the average length of molecules mapped to the reference genome.

### Validation by quantitative PCR

To perform quantitative PCR, we isolated genomic DNA by using a QIAamp DNA Mini Kit (QIAGEN). The frozen kidney tissue was ground in liquid nitrogen and subjected to DNA extraction *via the following protocol: DNA purification from tissues* (QIAamp DNA Mini Kit). For DNA isolation from sperm cells, DPBS buffer was added to frozen sperm pellets to a volume of 100 µL, followed by the procedure described in user-developed protocol 2: *Isolation of genomic DNA from sperm* (QA04; QIAamp DNA Mini Kit). The DNA concentration was measured with a QUBIT fluorometer via the Qubit dsDNA BR Assay Kit (Invitrogen).

Telomere length assessment was performed in triplicate for all the samples on a Bio-Rad CFX96 instrument. Two separate singleplex reactions were performed for each sample: one with telomere (T) primers (forward: 5’-CGG TTT GTT TGG GTT TGG GTT TGG GTT TGG GTT TGG GTT-3’; reverse: 5’-GGC TTG CCT TAC CCT TAC CCT TAC CCT TAC CCT TAC CCT-3’) and one with primers that amplify a part of the albumin gene (forward: 5’-CGG CGG CGG GCG CGG GCT GGG CGG CGG AAG TGC TGT AGT GGA TCC CTG-3’; reverse: 5’-GCC CGG CCC GCC GCG CCC GTC CCG CCG GAG AAG CAT GGC CGC CTT T-3’) as a single-copy reference (S). Reactions were performed in a 10 µl volume containing 5 µl of iTaq Universal SYBR Green Supermix (Bio-Rad), 20 ng of DNA, nuclease-free water, and forward and reverse primers at a final concentration of 0.3 µM each for the telomere target and 0.8 µM each for the albumin target. The samples were run in 96-well plates, and each plate contained three negative controls with nuclease-free water instead of DNA. The thermal profile for amplification of the albumin target was 10 min at 95 °C, followed by 40 cycles of 15 s at 95 °C, 10 s at 62 °C, and 15 s at 74 °C (data collection). For the telomere target, the thermal profile was 10 min at 95 °C, followed by 30 cycles of 15 s at 95 °C and 1 min at 56 °C (data collection). We used CFX Maestro 2.2 software (Bio-Rad) to extract the Ct values for telomere and albumin amplicons from each sample. Average Ct values from triplicate samples were used to calculate the relative telomere length (T/S ratio) according to the formula 2^-ΔΔCt^, where ΔΔCt = ΔCt(sample) – ΔCt(reference sample) and ΔCt = Ct(T) − Ct(S).

## Declarations

### Ethics approval

All procedures involving mice were approved by the Ministry of Agriculture, Forestry and Water Management of the Republic of Croatia, Class: UP/I-322-01/19-01/59 and case number: 525-10/0543-20-4, and carried out in compliance with relevant guidelines: the ILAR Guide for the Care and Use of Laboratory Animals, the EU Directive 2010/63/EU, the Croatian Animal Welfare Act (NN 102/17) and the Ruđer Bošković Institute’s policy on animal use and ethics.

### Consent for publication

Not applicable.

### Availability of data and materials

TelOMpy is written in the Python programming language. The package, along with all usage information, is available on the GitHub repository - https://github.com/ivanp1994/telompy. The OGM dataset used in the current study is available from the corresponding author upon reasonable request.

### Authors’ contributions

ŽP designed and led the research. IP developed the protocol for HMW DNA extraction from sperm cells, performed experiments, HMW DNA extractions, and PFGE. IP and ŽP developed TelOMpy, analyzed OGM data, and interpreted the results. IP wrote the code for TelOMpy. SS performed and analyzed the qPCR data. ŽP wrote the manuscript draft. All the authors reviewed the final manuscript.

## Acknowledgments

Data analysis was performed on the high-performance computing cluster at the University Computing Centre (SRCE), University of Zagreb. We thank Kristina Sporiš for technical support, Ranko Stojković for mouse facility support, Petra Mikolčević for help with preparing sperm cells, and Regina Bezděková Fillerová for help with optical genome mapping.

