## Supplementary for "TelOMpy enables single-molecule resolution of telomere length from optical genome mapping data"

**Table S1.** Concentrations of isolated HMW DNA after DLS labeling.

| Tissue | Group | Mouse (ID) | DNA Concentration* (ng/µL) | Coefficient of variation** (%) |
|---|---|---|---|---|
| **Sperm** | C | 5460 | 9.4 | 6.2 |
| **Sperm** | C | 5790 | 6.1 | 20.0 |
| **Sperm** | C | 5458 | 6.0 | 4.2 |
| **Sperm** | C | 5792 | 9.0 | 26.0 |
| **Sperm** | WD | 5455 | 8.7 | 5.1 |
| **Sperm** | WD | 5457 | 10.5 | 9.7 |
| **Sperm** | WD | 5456 | 6.8 | 2.1 |
| **Sperm** | WD | 5787 | 13.5 | 6.8 |
| **Sperm** | WD | 5788 | 8.3 | 11.7 |
| **Kidney** | C | 5460 | 12.0 | 9.8 |
| **Kidney** | C | 5458 | 3.2 | 0.2 |
| **Kidney** | C | 5790 | 7.1 | 0.1 |
| **Kidney** | C | 5792 | 6.6 | 0.2 |
| **Kidney** | WD | 5455 | 7.9 | 0.1 |
| **Kidney** | WD | 5456 | 4.8 | 0.0 |
| **Kidney** | WD | 5457 | 8.6 | 0.6 |
| **Kidney** | WD | 5787 | 5.9 | 0.2 |
| **Kidney** | WD | 5788 | 4.3 | 0.1 |

* Bionano’s recommendation: 4 - 16 ng/µL
** Bionano’s recommendation: < 30% between three measurements

**Table S2.** OGM data quality indicators

| Tissue | Group | Mouse (ID) | The average length of the molecules (filtered) / kbp | Label density / 100 kbp | Coverage of the reference before alignment / X | Fraction of molecules aligned | Effective coverage of reference / X | Diploid genome map N50 / Mbp |
|---|---|---|---|---|---|---|---|---|
| Kidney | C | 5458 | 258.3 | 15.6 | 273.1 | 0.9 | 192.7 | 96.0 |
| Kidney | C | 5460 | 275.4 | 15.3 | 764.6 | 0.9 | 561.9 | 108.0 |
| Kidney | C | 5790 | 288.0 | 15.8 | 657.1 | 0.9 | 542.6 | 111.2 |
| Kidney | C | 5792 | 255.8 | 15.8 | 598.3 | 0.9 | 472.0 | 118.0 |
| Kidney | WD | 5455 | 252.3 | 12.6 | 711.9 | 0.6 | 390.6 | 102.1 |
| Kidney | WD | 5456 | 276.5 | 16.4 | 463.3 | 0.9 | 356.0 | 108.8 |
| Kidney | WD | 5457 | 266.1 | 12.5 | 616.9 | 0.7 | 391.7 | 102.0 |
| Kidney | WD | 5787 | 253.1 | 16.0 | 574.3 | 0.9 | 459.2 | 102.0 |
| Kidney | WD | 5788 | 252.6 | 15.6 | 554.2 | 0.9 | 427.5 | 102.1 |
| Sperm | C | 5458 | 246.0 | 14.1 | 314.6 | 0.8 | 206.3 | 101.3 |
| Sperm | C | 5460 | 270.3 | 16.2 | 219.8 | 0.8 | 144.4 | 102.0 |
| Sperm | C | 5790 | 282.9 | 14.1 | 264.2 | 0.8 | 188.0 | 106.1 |
| Sperm | C | 5792 | 269.9 | 14.3 | 288.2 | 0.8 | 195.3 | 96.4 |
| Sperm | WD | 5455 | 265.2 | 15.7 | 359.8 | 0.8 | 230.2 | 102.0 |
| Sperm | WD | 5456 | 259.0 | 14.7 | 232.7 | 0.8 | 171.4 | 95.0 |
| Sperm | WD | 5457 | 240.3 | 16.1 | 117.3 | 0.9 | 79.6 | 96.1 |
| Sperm | WD | 5787 | 252.2 | 15.0 | 470.4 | 0.9 | 320.6 | 95.9 |
| Sperm | WD | 5788 | 264.6 | 13.9 | 313.7 | 0.8 | 196.7 | 101.2 |
| **Bionano recommendation (target values):** | | | **230** | **14 - 17** | **> 100** | **> 0.6** | **> 70** | **> 50** |

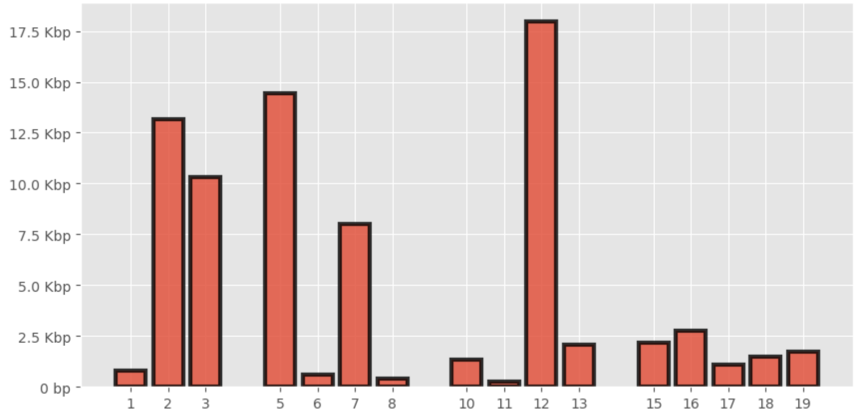


**Figure S1.** Distance between the last DLE-1 motif and the annotated start of the telomere on the q arm of the mouse reference genome (mm10). Data are presented for chromosome arms (x-axis) suitable for the analysis in this study.

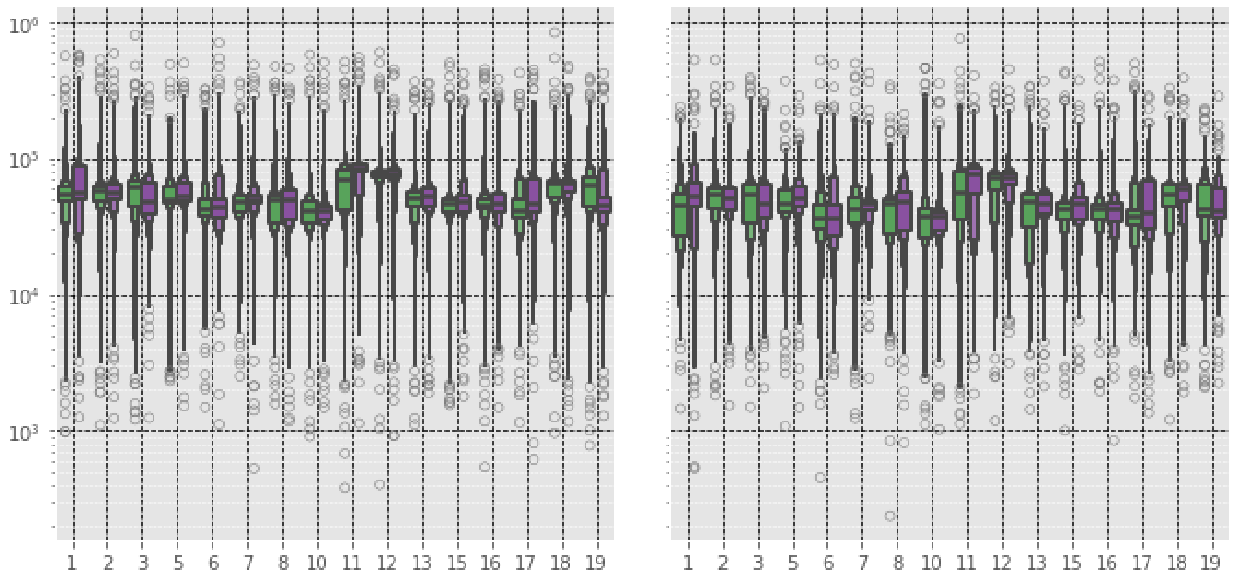


**Figure S2.** Distribution of telomere lengths of mice from the experimental (green) and control (purple) groups by chromosomes shown by boxen plot for kidney (left) and sperm (right) samples. The horizontal line in the widest part of each of the distributions represents the median length.

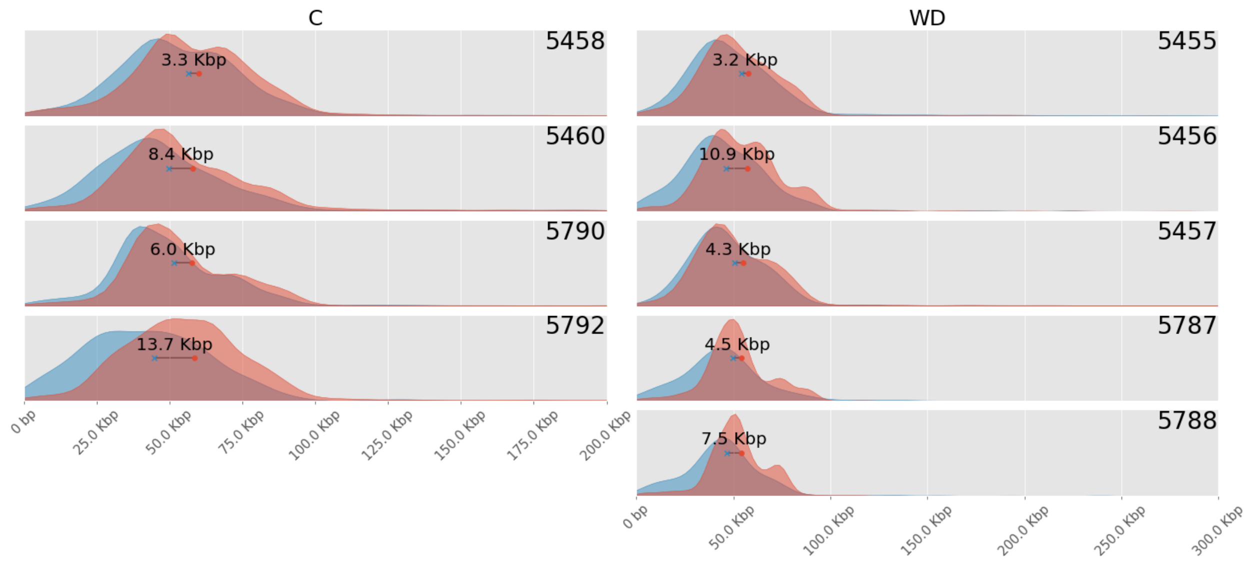


**Figure S3.** Comparison of telomere length distributions between sperm (blue) and kidney (orange). Comparisons are shown for individual mice from the control (left) and experimental (right) groups. The difference in average telomere length is indicated on each graph.

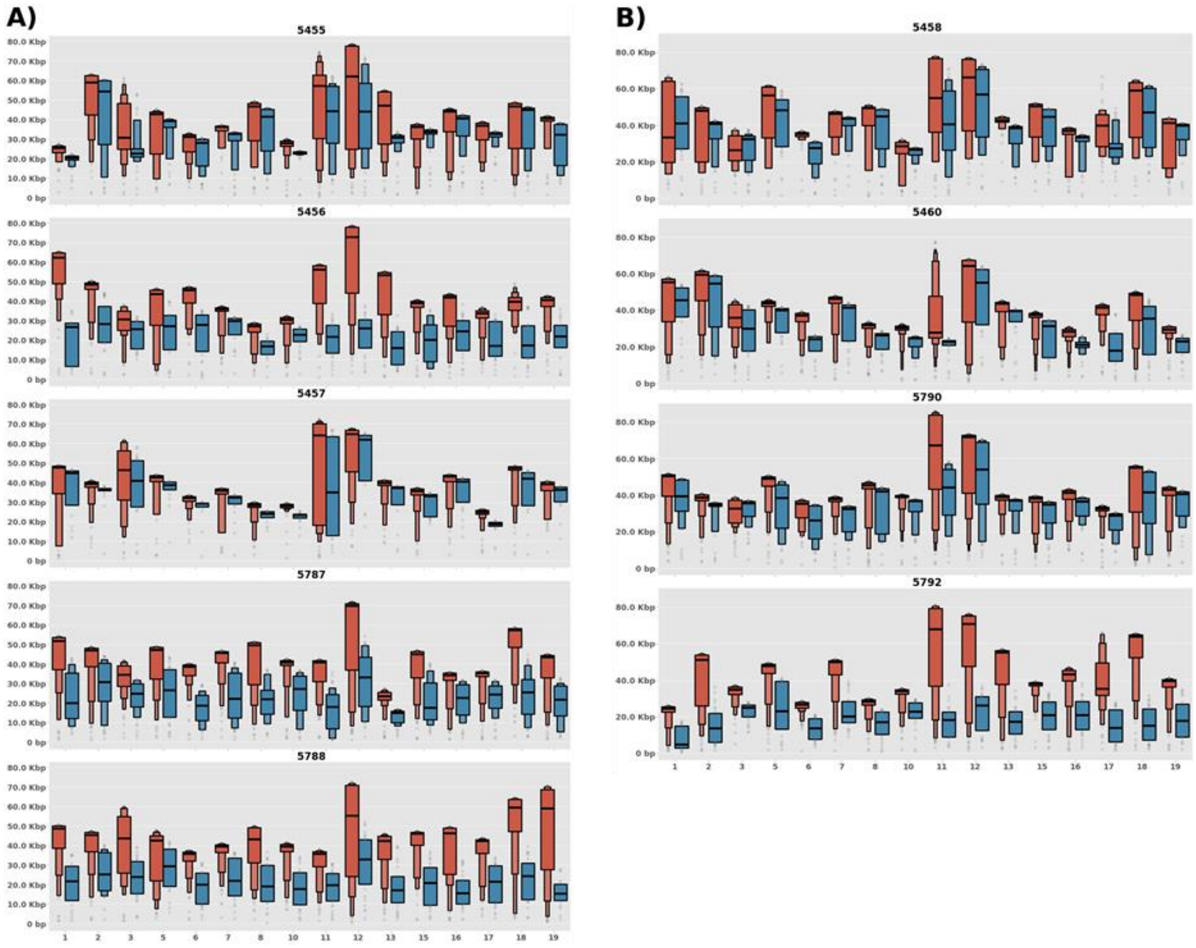


**Figure S4.** Distribution of telomere length in the Q1 per chromosome. Data for individual mice from the experimental (A) and control (B) groups are shown for sperm (blue) and kidneys (orange). The mouse identification number is indicated above each plot.

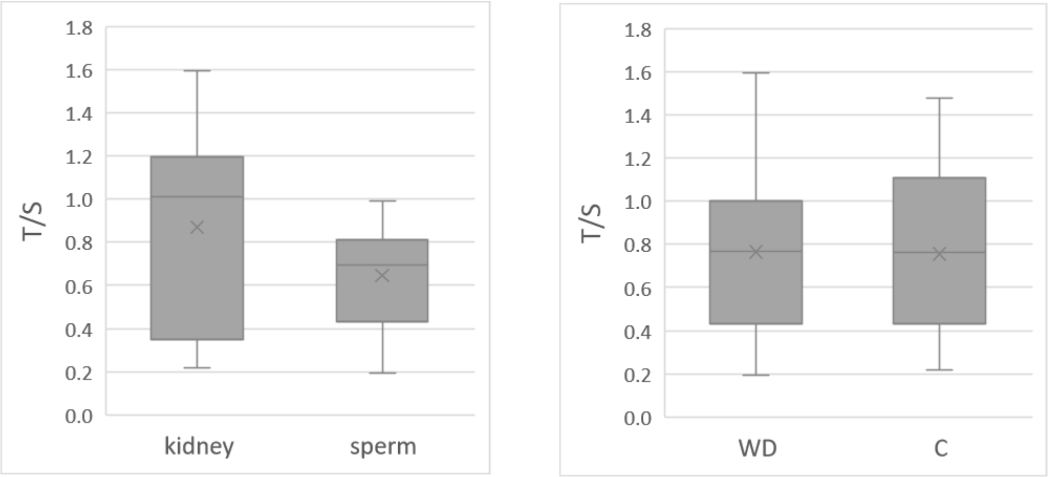


**Figure S5.** qPCR on an extended set of samples in the high-fat diet experiment. Telomere lengths relative to a control sample are presented as T/S ratio in comparison between tissues (left) and groups (right). N(kidney) = 20; N(sperm) = 19; N(WD) = 20; N(C) = 19. Results suggest the shorter telomeres in sperm compared to kidney (two-tailed paired t-test; p-val = $6.3 \times 10^{-3}$) and no significant difference in telomere length between the control and experimental group (two-tailed unpaired t-test; pval = 0.892).

**Text S1.** Protocol for isolation of HMW DNA from sperm

### Protocol for isolation of HMW DNA from mouse sperm

(Compliant with optical genome mapping standards)

The composition of solutions and buffers is given in Table S3.

Chemicals and reagents are listed in Table S4.

#### Preparation of agarose plugs

1. Low melting point agarose (LMP) was weighed and suspended in DPBS buffer to a final mass (w/v) concentration of 2% (2 mg of LMP agarose per 100 µL of DPBS). Agarose was dissolved in a boiling water bath and transferred to a water bath at a temperature of 45 °C to cool down.
2. Pellets of sperm cells (stored at -80 °C) were suspended in DPBS to 3-4 million cells in a 40 µL volume. The suspension was transferred to a water bath at a temperature of 45 °C.
3. After a short time, the sperm suspension was mixed with a cooled-down LMP agarose solution and resuspended several times with a pipette for even mixing. 80 µL was aliquoted into BioRad mold(s), corresponding to 3-4 million sperm cells per agarose plug. The molds were kept at 2-8 °C for 20-30 minutes until gel was formed.

#### Cell lysis and washing

1. After the agarose plug solidified, they were transferred to Eppendorf tubes (one agarose block per tube), and 500 µL of Lysis Buffer was added, corresponding to ~10 times the plug volume of the plug.
2. Eppendorf tubes containing agarose plug in Lysis Buffer were placed horizontally in a hybridization oven and incubated for 16 hours at 50 °C (working temperature of proteinase K) at 3 rpm.
3. After the 16-hour incubation, another 50 µL of proteinase K (concentration 20 mg/mL, i.e., 1 mg per block) was added, and the mixture was incubated for an additional hour under the same conditions.
4. Agarose plugs were washed for 30 min on ice with ~10 times their volume of each of the indicated buffers, using minimal agitation on an orbital shaker (100 rpm):
   - Once with ice-cold 50 mM EDTA (pH 8)
   - Once with ice-cold TE 1 X (pH 8)
   - Three times with ice-cold TE 1 X (pH 8) 0.1 mM PMSF (1 µL of 100 mM PMSF solution in isopropanol per 1 mL of buffer TE 1 X)
   - Three times with ice TE 1 X (pH 8)
5. After the last wash, agarose plugs were transferred to Wash Buffer and stored at 2-8 °C.

#### Restriction enzyme digestion

1. Half of the agarose plug (~40 µL volume) was transferred to an Eppendorf tube containing 100 µL of 1X EcoRI buffer and incubated for 15 minutes at room temperature.
2. Agarose plug was then removed from the tube and transferred to a new Eppendorf tube containing 100 µL of 1X EcoRI buffer with 20 U of EcoRI restriction enzyme, and the mixture was incubated for 3-4 hours at 37°C.

#### Gel electrophoresis in a pulsed field

1. A 1% agarose gel was prepared by suspending 1.5 g of agarose per 150 mL 0.5X TBE buffer in an Erlenmeyer flask. The flask was covered with a piece of paper and heated in the microwave until the boiling point, after which it was removed from the microwave and gently stirred. The procedure was repeated until the solution became transparent. The flask was placed in a water bath heated to 50 °C for 15 to 20 minutes to cool down. The agarose solution was then poured onto the casting stand and left at room temperature for 30 minutes to solidify.
2. Prepared DNA agarose plugs were removed from the buffer and placed on a clean piece of parafilm, and excess buffer was removed by pipetting. Agarose plugs were inserted into wells on agarose gel. Lambda PFG Ladder was placed into the side wells. Wells were sealed with a 1% LMP agarose solution cooled down to 50 °C, and the gel was left at room temperature for 10 to 15 minutes.
3. The CHEF-DR® III Pulsed Field Electrophoresis System (Bio-Rad) was washed with 2 L of distilled water. About 2.5 L of 0.5 X TBE buffer was added, and buffer circulation was started with the cooling system.
4. When the temperature reached 14 °C, the gel with the samples was placed in the central chamber of the CHEF system. Electrophoresis was run under the following settings: switch time 5/50 s, run time 16 hrs, 6 Volts/cm, included angle 120°, temperature 14 °C.
5. The gel was stained by immersion in ~0.5 L 0.5X TBE buffer with 1 µg/mL ethidium bromide. After approximately 30 min, it was removed from the solution and photographed on a gel imaging system.

**Table S3.** List of buffers and solutions

| BUFFER | pH | COMPOSITION | PREPARATION |
|---|---|---|---|
| **EDTA** | 8 | 1.0 M EDTA | 372.24 g of EDTA (MW=372.24) was added to ~800 mL of redistilled water. The suspension was transferred to a magnetic stirrer/heater, and NaOH pellets were gradually added until the solution became clear. The volume was made up to ~980 mL with reH20. The pH was adjusted to 8, volume adjusted to 1 L, autoclaved |
| **Tris-HCL** | 8 | 1.0 M Tris-HCl | 24.23 g Trizma™ base added to 160 mL redistilled water. pH adjusted to 8 with HCl (37 %, 12 M), made up to 200 mL, autoclaved |
| **TBE 5x** | ~8.3 | 450 mM Trizma™<br>450 mM boric acid<br>10 mM EDTA | 27.82g boric acid (MW=61.83), 54.51g Trizma™ base (MW=121.14) and 20mL 0.5M EDTA (pH=8) in ~800 mL redistilled water, mixed on magnetic stirrer until dissolved. Adjusted volume to 1 L, autoclaved |
| **TE 1x** | 8 | 10 mM Tris-HCl,<br>1 mM EDTA | 10 mL 1.0 M Tris-HCl and 2 mL 0.5 M EDTA in ~980 mL redistilled water. pH adjusted to 8, volume adjusted to 1L, autoclaved |
| **Wash Buffer** | 8 | 10 mM Tris-HCl,<br>50 mM EDTA, | 18.61g EDTA (M2=372.24) i 1.21 g Trizma™ base in ~980mL redistilled water. pH adjusted 8, volume adjusted to 1L, autoclaved |
| **TBE 0.5x / EtBr** | ~8.3 | TBE 0.5X,<br>Ethidium bromide 1 mg/L | 100 mL TBE 5X diluted to 1 L with redistilled water. Added 100 µL solution of 10m g/mL ethidium bromide |
| **Sarkosyl™** | - | 25 % Sarkosyl™ | 5 g Sarkosyl™ powder added to ~12 mL redistilled water and mixed while heating. Volume adjusted to 20 mL, autoclaved |
| **Proteinase K** | - | 20 mg/mL Proteinase K | 10 mg proteinase K dissolved in 0.5 mL nuclease-free water |
| **DTT** | - | 1.0 M DTT | 154.25 mg DTT powder dissolved in 1 mL nuclease-free water |
| **Lysis Buffer** | 8 | 350 mM EDTA<br>2 % (w/v) Sarkosyl™<br>2 mg/mL Proteinase K<br>80 mM DTT | Prepare fresh. Mix 4.2 mL 0.5 M EDTA, 0.6mL 25 % Sarkosyl™, 0.6 mL 20mg/mL Proteinase K, 0.48 mL 1 M DTT. Adjust to 6 mL with nuclease-free water |

**Table S4.** List of chemicals and reagents

| CHEMICALS AND REAGENTS | MANUFACTURER AND CATALOG NUMBER |
|---|---|
| EDTA (powder) | Gram Mol P134122 |
| Trizma™ (powder) | Sigma Aldrich T1503 |
| Boric acid (powder) | Kemika EC-br. 233-139-2 |
| NaOH (pellets) | Kemika EC-br. 215-185-5 |
| HCl (37 %, 12 M) | Kemika EC-br. 231-595-7 |
| DPBS buffer 1x (solution) | Sigma Aldrich D8537 |
| EDTA (solution 0.5 M pH 8) | Sigma Aldrich E7889 |
| Proteinase K (powder) | Sigma Aldrich P2308 |
| Sarkosyl™ (powder) | Sigma Aldrich L-5125 |
| DTT (powder) | Thermo Scientific Catalog No. R0861 |
| Nuclease-free water | Ambion AM9937 |
| PMSF (solution, 0.1 M in isopropyl alcohol) | Boston BioProducts, SKU#: BP-481 |
| LMP Agarose | Cambrex BioScience InCert™ Part No. 50121 |
| Agarose | Bio-Rad Catalog 162-0137 |
| EcoR1 (20000 U/mL) | NEB R0101S |
| EcoR1 Buffer (10x) | NEB R0101S |
| Lambda PFG Ladder | NEB N0341S |
| Ethidium bromide (solution 10m g/mL) | Promega, H5041 |
